# Generating a Collection of Single-Gene Deletions in the *eps* Operon in *Bacillus subtilis* Strain NCIB 3610

**DOI:** 10.64898/2026.10.05.756831

**Authors:** Nicole T. Cavanaugh, Bryanna P. Upton, Eva V. Franco, Leticia Lima Angelini, Cameron W. Habib, Juliana Xu, Claire Chai, Emily Wu, Maor Bar-Peled, Yunrong Chai

**Affiliations:** Northeastern University, College of Science, Department of Biology, Boston, MA 02115; Northeastern University, Bouvé College of Health Sciences, Boston, MA 02115; Northeastern University, College of Science, Department of Psychology, Boston, MA 02115; Brookline High School, Brookline, MA 02445; University of Georgia, Complex Carbohydrate Research Center, Athens, GA 30602

**Author notes:** Contributions: Nicole Cavanaugh: strain construction, primer design, biofilm growth and imaging, writing, figures Bryanna Upton: strain construction, biofilm growth and imaging, figures, writing Leticia Lima Angelini, Claire Chai: strain construction Cameron Habib: strain procurement from BGSC, primer design Eva Franco, Juliana Xu, Emily Wu: strain construction, biofilm growth and imaging.

## Abstract

In the environment, bacteria mainly exist in multicellular, surface-adhered communities called biofilms. One key feature of the biofilms is a self-produced extracellular matrix that embeds cells within a structured community and contributes to their adhesion, organization, and retention. The matrix of most characterized biofilms contains three primary components: polysaccharides, protein, and extracellular DNA (eDNA). Upon biofilm induction, *Bacillus subtilis* produces and secretes a polysaccharide known as exopolysaccharide (EPS). The 15 genes of the *epsA-O* operon encode proteins that coordinate the assembly, export, and polymerization of EPS oligosaccharide (glycan) subunits. While much progress has been made to elucidate the chemical structure of EPS, further work is needed to determine the contributions of some *eps* genes to EPS biosynthesis experimentally. Here, we generated a collection of marker-less, in-frame deletions targeting each *eps* gene in the *epsA-O* operon in *B. subtilis* strain NCIB 3610 using the Cre system. Biofilm phenotypes of resulting mutants were used to assess each gene’s importance to biofilm development.

**Impact:** The *epsA-O* single-gene deletion collection was designed to enable systematic assessment of the contribution of each *eps* gene to EPS production and biofilm-associated phenotypes. When combined with current knowledge of EPS structure, this collection will help clarify the roles of individual genes in EPS biosynthesis. This approach can also be easily adapted for the systematic analyses of other large, complex operons in diverse bacterial species.

## Introduction

### Biofilm is a multicellular community of microorganisms

In nature, microorganisms form multicellular communities called biofilms. These communities are embedded in a self-produced extracellular matrix, composed mainly of polysaccharides, proteins, and extracellular DNA (eDNA), which contribute to cell-cell cohesion and adhesion to surfaces [1]. This matrix provides many benefits to the community including providing a barrier to antibiotic penetration, protection from UV radiation, and defense against nearby antagonistic microorganisms [2]. It also provides a medium through which individual cells can move and communicate via chemical signals. Additionally, cells within the community can differentiate into specialized cells with distinct physiological properties, allowing different subpopulations to contribute to matrix production, motility, stress resistance, and other functions essential for biofilm development. Biofilms are thus important for bacterial survival because they promote surface colonization, environmental persistence, and resistance to diverse stresses.

### *Bacillus subtilis* as a model organism to study biofilm formation

*Bacillus subtilis* is a Gram-positive, non-pathogenic, rod-shaped soil bacterium [3], [4]. Under laboratory settings, *B. subtilis* forms two types of biofilms: colony biofilms, which form on a solid surface, and pellicle biofilms, which form at the air-liquid interface [5]. These biofilms are architecturally robust and morphologically distinct, contributing to the use of *B. subtilis* as a model organism for biofilm studies. In nature, *B. subtilis* also forms beneficial biofilms on plant roots [6], [7]. These biofilms help protect the plant from pathogenic microorganisms in the soil, while the plant provides sugars and nutrients to the bacteria.

In *B. subtilis,* the extracellular matrix of the biofilm primarily consists of proteins and polysaccharides. Three main extracellular proteins contribute to matrix formation. The first two are TasA and TapA. TasA is a matrix protein that forms amyloid-like fibers. TapA anchors and facilitates TasA fiber formation. Both proteins are encoded in the *tapA-sipA-tasA* operon [8], [9]. The third protein is BslA, which is a self-assembling hydrophobin that forms a water-repellent surface layer on the biofilm [10]. The main polysaccharide produced during *B. subtilis* biofilm formation is an exopolysaccharide (EPS), which is synthesized via a 15-gene *epsA-O* operon and functions as a “glue” which contributes to cell-cell cohesion within the biofilm [11].

### EPS Biosynthesis in *B. subtilis*

In the *B. subtilis* biofilm model strain NCIB 3610, EPS is the only currently identified polysaccharide required for biofilm formation. It is a hydrated polysaccharide that retains moisture for the biofilm and allows for capture of nutrients from the surrounding environment while aiding in biofilm adhesion, cell aggregation, cohesion, and biofilm protection from environmental stresses [12]. The EPS biosynthesis pathway in *B. subtilis* is highly complex, requiring the assembly of six sugars into a repeating glycan unit within the cytosol using a membrane-bound lipid carrier, followed by its export across the cell membrane and polymerization [13]. Many well-established systems for studying complex exopolysaccharide biosynthesis are found in Gram-negative models, including the *pgl* operon in *Campylobacter jejuni* and the LPS biosynthetic pathway in *E. coli* [14], [15], [16]. The *pgl* operon in *Campylobacter jejuni* has been especially informative due to its similarity in function to the *eps* operon in *B. subtilis* [17]. Some Gram-positive models include the *cap* operon in *Bacillus cereus* and *Bacillus anthracis*, each consisting of five genes that work together to produce extracellular capsules [18], [19].

The *eps* operon (*epsA-O*) contains 15 genes, whose encoded proteins work together to assemble, export, and polymerize EPS subunits [20](Figure 1). The operon encodes six glycosyl-transferases (GTs), including the initiating EpsL [17], a phosphoglycosyl-transferase (PGT) that transfers a phosphor-sugar from UDP-sugar donor to undecaprenyl - phosphate (Und-P, C55-P), a membrane-bound lipid carrier. In an *in vitro* assay, the sugar transferred by EpsL was determined to be *N,N*′-diacetylbacillosamine (diNAcBac, also referred to as Qui*p*NAc4NAc), yielding Und-PP-Qui*p*NAc4NAc [17]. The preceding GT,

**Figure 1:**
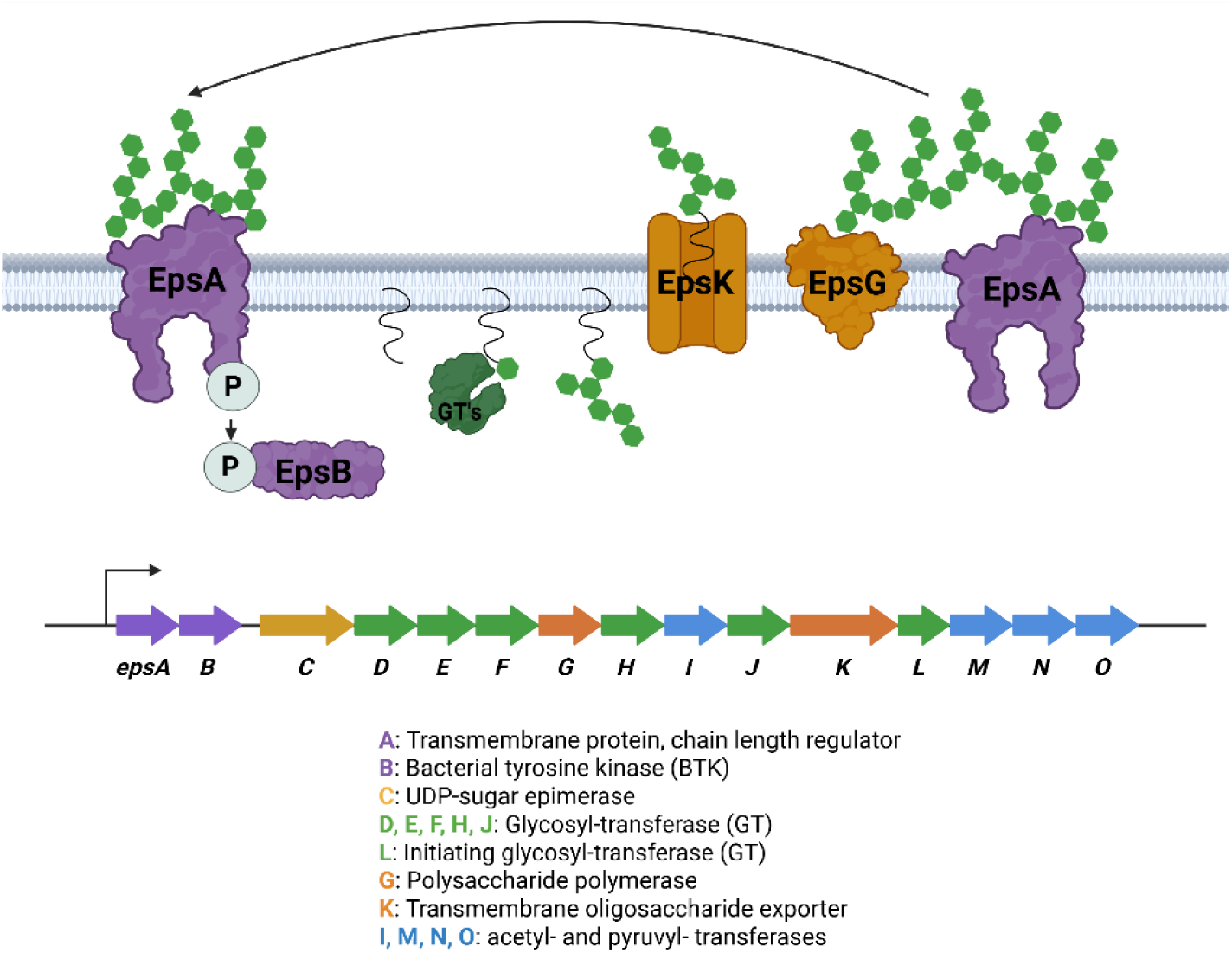
The *eps* operon and predicted biosynthesis pathway. The *eps* operon consists of 15 genes involved in EPS assembly. Glycosyltransferases, shown in green, take turns adding building block sugars onto a membrane-bound lipid carrier. Once the fully formed EPS glycan is complete, it is exported through the EpsK flippase and polymerized into long chains by EpsG, both enzymes are shown in orange. The length of those chains is mediated by EpsA-EpsB, shown in purple, and when the desired length is formed, the sugar is released into the matrix. Synthesized EPS molecules are also capable of binding to EpsA, prompting a signaling cascade that impacts the regulation of EPS biosynthesis.

EpsD, adds the next sugar onto the growing glycan chain [17]. Its function was also determined *in vitro*, finding that EpsD uses UDP-GalNAc as a sugar donor and transfers the GalNAc moiety, generating C55-PP-QuiNAc4NAc-GalNAc. These are the only two GTs, as of now, that have been determined biochemically. [17].

The chemical structure of EPS was recently elucidated [13]. According to nuclear magnetic resonance (NMR) analysis of EPS purified from biofilms formed by *B. subtilis* strain NCIB 3610, a single EPS building block glycan consists of a three-sugar backbone and a three-sugar branch attached to the center backbone sugar. The backbone consists of Qui*p*NAc4NAc, Gal*p*NAc, and Glc*p*NAc, listed in order from the Qui*p*NAc4NAc attached to the lipid carrier [13]. The identity of the second backbone sugar differs slightly from the one identified and proposed by Arbor et al. in 2023. That study reported that EpsD preferentially adds glucose to the growing glycan chain, whereas galactose was incorporated with substantially lower efficiency [17]. The branching sugar consists of a chain of three galactose residues extending from the central backbone sugar [13].

EpsE, EpsF, EpsH, and EpsJ are predicted to transfer distinct sugars from UDP-sugar substrate to the growing glycan chain on the lipid carrier. The UDP-sugar substrate recognized by each GT has not yet been experimentally determined. However, some predictions can be made based on protein sequence homology. EpsF most likely adds the third sugar, glucose, onto the EPS backbone to forming an α(1→3 ) linkage between galactose and glucose [13]. EpsE most likely adds the first galactose onto the branching sugar, forming an α(1→4) linkage between the backbone and first branching galactose molecules. The final two GT’s, EpsH and EpsJ, both are predicted to form β(1→6) linkage between branching galactose molecules. Their structural similarity makes it difficult to determine their respective functional differences.

The *eps* operon also encodes a few genes whose functions are involved in substrate sugar modification. For instance, EpsC is a UDP-sugar epimerase, which can invert the stereochemistry of sugar molecules to create the Qui*p*NAc needed for EPS formation [21], [22]. EpsM and EpsN are putative acetyl-transferases, which add acetyl groups to the backbone sugars [23], [24]. EpsI and EpsO are predicted, based on bioinformatic analyses, to be pyruvyl-transferases, likely involved in pyruvylation of the last two branching galactoses on EPS (Figure 5). However, which galactose residue each of them pyruvates is unknown.

**Figure 5:**
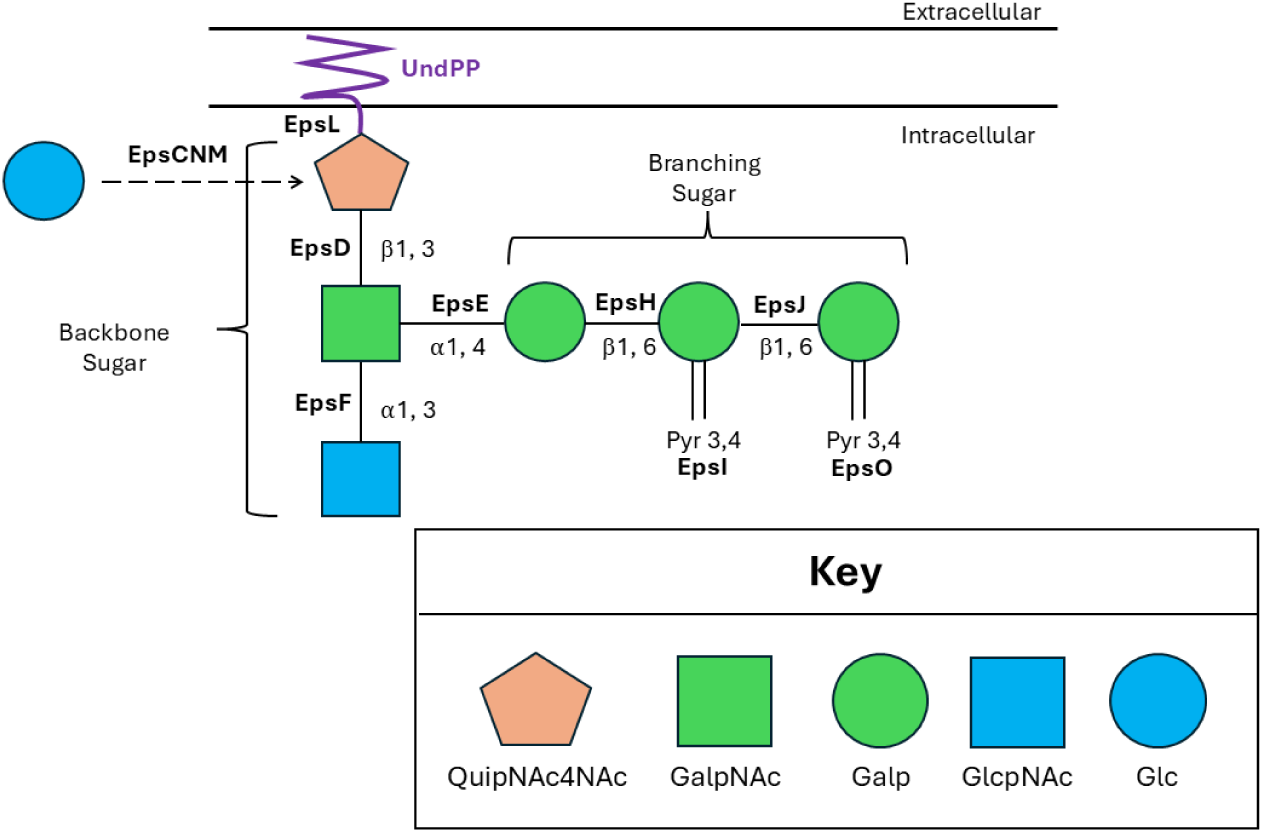
Proposed hypothetical model for biosynthesis of EPS molecules. In this model, EpsL attaches Qui*p*NAc4NAc to a lipid carrier attached to the membrane to initiate the building of EPS. This is followed by the addition of GalpNAc by EpsD and GlcpNAc by EpsF. EpsE adds the first galactose to form the branching sugar. Afterwards EpsH and EpsJ add the second and third galactose molecules to the side chain, respectively. Additionally, EpsI may pyruvylate the second branching galactose while EpsO may pyruvylate the last branching galactose.

It might be worth pointing out that most UDP-sugar substrates for EPS assembly, such as UDP-glucose and UDP-galactose, are synthesized by enzymes outside the operon. For example, UDP-glucose is generated from the glycolytic intermediate glucose-6-phosphate via two key enzymes: PgcA (phosphoglucomutase), which converts glucose-6-phosphate to glucose-1-phosphate, and GtaB (UTP-glucose-1-phosphate uridylyltransferase), which converts glucose-1-phosphate into UDP-glucose utilizing UTP [25]. This UDP-glucose can be further converted into UDP-galactose by the UDP-glucose-4-epimerase GalE, which also mediates the reversible conversion between UDP-N-acetylgalactosamine and UDP-N-acetylglucosamine [26], [27]. Additionally, when environmental galactose is available, UDP-galactose can be synthesized through the Leloir pathway via GalK (galactose kinase), GalT (galactose-1-phosphate uridylyltransferase), and GalE [26], [28].

Once the complete six sugar glycan repeat is assembled on the membrane-linked lipid, the resulting C55-PP-glycan is flipped from the cytoplasmic side to the extracellular side of the membrane by a predicted flippase, EpsK [29]. The glycan repeats are then polymerized by the predicted EpsG polymerase to form mature EPS chains [30]. Two additional genes in the operon, *epsA* and *epsB*, encode putative chain-length determination proteins [31]. The mechanism of EPS polymerization and chain length control remained poorly understood in *B. subtilis*. EpsG is homologous to the *E. coli* Wzy polymerase involved in lipopolysaccharide biosynthesis. EpsA and EpsB, based on some homology, resemble the better characterized *E. coli* Wzz chain-length regulator. In *E. coli*, Wzz is one single transmembrane chain-length regulator with an extracellular domain that is proposed to interact with components of the polymerization machinery [14]. The duration and stability of these interactions, together with structural constrains imposed by Wzz, are thought to influence LPS chain length [14]. In *B. subtilis*, EpsB is a cytoplasmic bacterial tyrosine kinase (BTK) and resembles the C-terminal domain of Wzz, while EpsA is a transmembrane protein which resembles the N-terminal domain of Wzz and acts as a BTK modulator [32]. It was recently predicted using AlphaFold-Multimer that EpsA and EpsB may form a complex [33].

While substantial stride has been made to elucidate the functions of each *eps* gene in recent years, only two GT functions (EpsL and EpsD) have been confirmed experimentally [17]. Roles of other GTs and sugar modification enzymes remain inferred from sequence homology to genes in other organisms and from the known chemical structure of EPS. We created a collection of marker-less, in-frame deletions of each *eps* gene in a biofilm-forming strain of *B. subtilis*. Through biofilm assays, we determined how the absence of each gene impacted biofilm physiology. These changes in biofilm development, along with current knowledge of EPS structure, give us important insight into the importance of each gene product which can help direct future chemical studies using NMR. This method is able to be adapted to large, complex operons in species important to the development of biofilms or other extracellular polysaccharides.

## Methods

### Construction of insertional gene deletion of *eps* genes in NCIB 3610

Strains with insertional deletions of each *eps* gene in *B. subtilis* strain 168 were obtained from the *Bacillus* Genetic Stock Center (BGSC) (Table 1). Strains from the BGSC include an erythromycin resistance cassette inserted into the appropriate gene site flanked by *lox* sites. These *lox* sites allow further creation of marker-less, in-frame gene deletions [34].

**Table 1:** Strains from BGSC.

| BGSC Number | Genotype | Host Strain | Resistance |
| --- | --- | --- | --- |
| BKE34370 | *epsA::erm* | 168 | MLS |
| BKE34360 | *epsB::erm* | 168 | MLS |
| BKE34350 | *epsC::erm* | 168 | MLS |
| BKE34340 | *epsD::erm* | 168 | MLS |
| BKE34330 | *epsE::erm* | 168 | MLS |
| BKE34320 | *epsF::erm* | 168 | MLS |
| BKE34310 | *epsG::erm* | 168 | MLS |
| BKE34300 | *epsH::erm* | 168 | MLS |
| BKE34290 | *epsI::erm* | 168 | MLS |
| BKE34280 | *epsJ::erm* | 168 | MLS |
| BKE34265 | *epsK::erm* | 168 | MLS |
| BKE34250 | *epsL::erm* | 168 | MLS |
| BKE34240 | *epsM::erm* | 168 | MLS |
| BKE34230 | *epsN::erm* | 168 | MLS |
| BKE34220 | *epsO::erm* | 168 | MLS |
| ECE274 | pDR244 | *E. coli* | Amp |

To isolate the genomic DNA (gDNA) from each BGSC-sourced strain, a single colony of bacteria was grown in a 2 mL culture of LB broth supplemented with 1X MLS (24 µg/mL lincomycin and 1 µg/ml erythromycin). gDNA was purified using the Wizard Genomic DNA Purification Kit (Promega, USA). To introduce the insertional gene deletion into strain NCIB 3610, NCIB 3610 was grown in 2 ml of MC medium for four hours shaking at 37°C. After four hours of incubation, 300 µL of culture was mixed with 10 µL of purified gDNA from a selected 168 strain. After an additional two hours of growth at 37°C, the 300 µL cells and gDNA mixture was spread on an LB agar plate containing 1X MLS and grown at 37°C overnight. The following day, colonies of MLS-resistant transformants were picked and streaked on fresh LB+MLS plates to obtain single colonies.

### Construction of marker-less in-frame gene deletion in NCIB 3610

Erythromycin-resistance cassettes and the associated *lox* sites were removed from the chromosome of the *eps* insertional gene-deletion mutant strains of NCIB 3610 using the loop-out plasmid, pDR244 [34](Figure 2). pDR244 harbors the Cre-recombinase gene, encoding a site-specific recombinase that recognizes the *lox* sites flanking the erythromycin-resistance cassette and catalyzes recombination between them, thereby excising the antibiotic cassette from the chromosome. To prepare the plasmid, 2 mL LB culture of ECE274 (Table 1) supplemented with 100 µg/mL ampicillin was grown at 30°C for five hours. The plasmid was purified using the GeneJET Plasmid Miniprep Kit (Thermo Fisher Scientific, Waltham, MA, USA) according to the manufacturer’s instructions. Each insertional gene-deletion strain in NCIB 3610 was grown in 2 mL of competence medium (MC) for four hours with shaking at 37°C. After four hours, 300 µL of culture was mixed with 10 µl of pDR244 and grown for an additional two more hours with shaking at 37°C to facilitate transformation. The entire transformation mixture (300 µL) was plated on LB agar plates containing 1X MLS. Plates were incubated at 30°C overnight.

**Figure 2:**
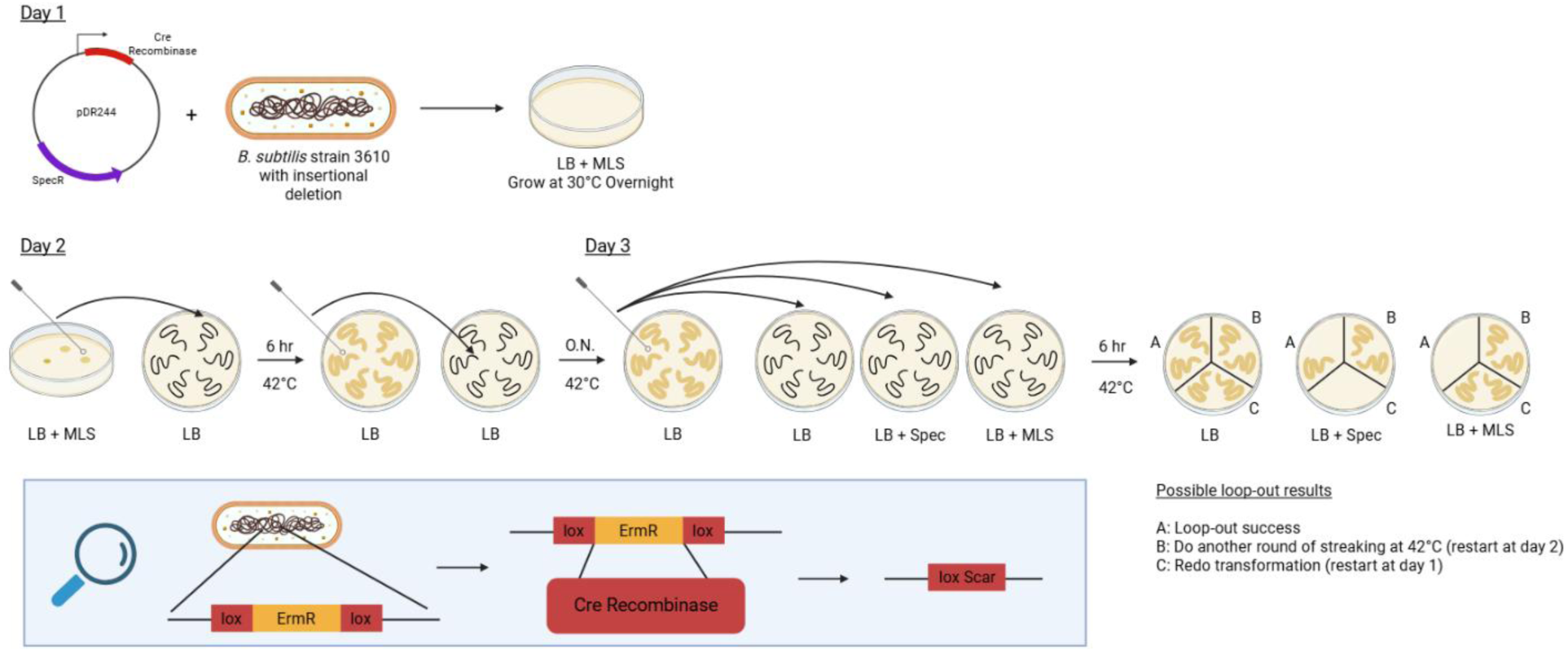
Creating non-polar deletions using the Cre loop-out system.

Several successful transformants from each strain were streaked onto a fresh LB plate and incubated at 42°C, the non-permissive temperature for replication of the temperature-sensitive pDR244 plasmid [35]. After six hours of growth, each transformant was restreaked onto a fresh LB plate and incubated overnight at 42°C. This incubation was used to promote loss of pDR244 after Cre-mediated excision of the antibiotic cassette. To screen for successful cassette excision and plasmid loss, each transformant was streaked onto three different agar plates, LB, LB with 100 µg/mL spectinomycin, and LB with 1X MLS, and all plates were incubated at 42°C for six hours. Because pDR244 confers spectinomycin resistance, strains in which Cre-mediated excision had occurred were expected to grow on LB but not LB with 1X MLS. Therefore, candidate marker-less mutants were identified by growth on LB and the absence of growth on both spectinomycin- and MLS-containing plates.

Transformants that grew on spectinomycin were presumed to retain pDR244 and were subjected to additional rounds of restreaking at 42°C to promote plasmid cure. Transformants that did not grow on spectinomycin but continued to grow on MLS had likely loss of pDR244 without undergoing the desired Cre-mediated excision; these strains were therefore retransformed with pDR244.

Candidate marker-less mutants were confirmed by colony PCR using primers that flank the deleted region. The PCR products were analyzed for expected size change and verified by DNA sequencing to confirm precise in-frame gene deletion. Growth on the appropriate selective agar plates were recorded for each confirmed mutant.

Successful marker-less gene deletions were confirmed by PCR using OneTaq Quick Load 2x Master Mix (New England Biolabs, Ipswich, MA, USA 01938). Primers used are outlined in Table 2.

**Table 2:**
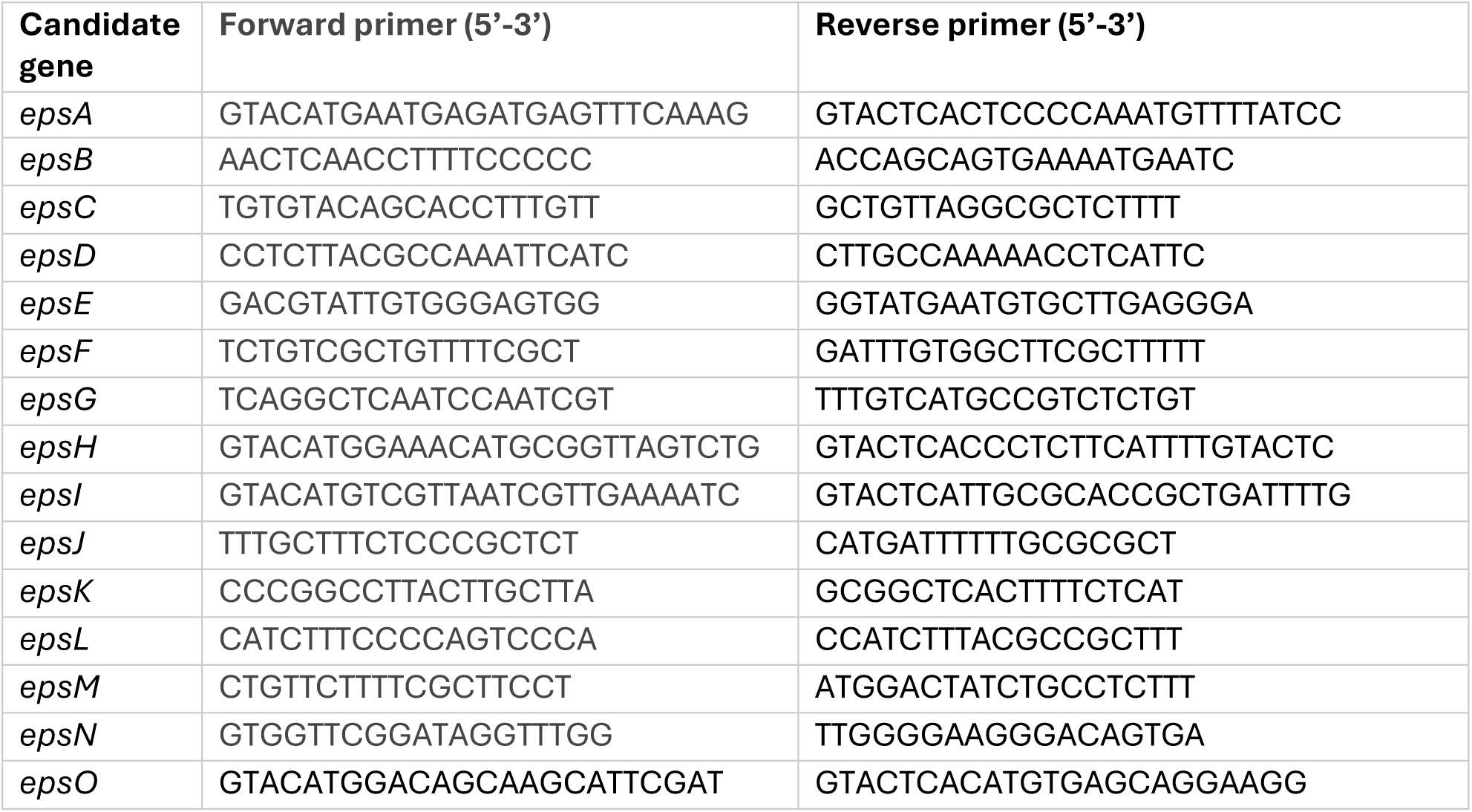
Primers to confirm marker-less gene deletion in NCIB 3610.

### Vector construction for genetic complementation

A custom vector was generated to facilitate the complementation of individual *eps* gene deletion by expressing each gene under the control of the by the *epsA* promoter (P*_epsA_*) at the chromosomal *amyE* locus (Figure 3). The *epsA* promoter (P*_epsA_*) was amplified from NCIB 3610 gDNA using primers FP (5’-GTC<u>GAATTC</u>CTAGAAATTCTCCTCTATTCCTGTCG-3’), RP (5’-GATC<u>AAGCTT</u>CATAGCCTTCAGCCTTCC-3’), and Gloria Nova HS 2X Master Mix (ABclonal, Woburn, MA, USA, 01801). The amplified prompter DNA fragment was designed to contain EcoRI and HindIII compatible ends and was cloned between the EcoRI and HindIII restriction sites of pDG1662 to generate pYC234 (restriction enzymes from New England Biolabs, Ipswich, MA, USA 01938). Ligation was performed using T4 DNA Ligase (ABclonal, Woburn, MA, USA, 01801).

**Figure 3:**
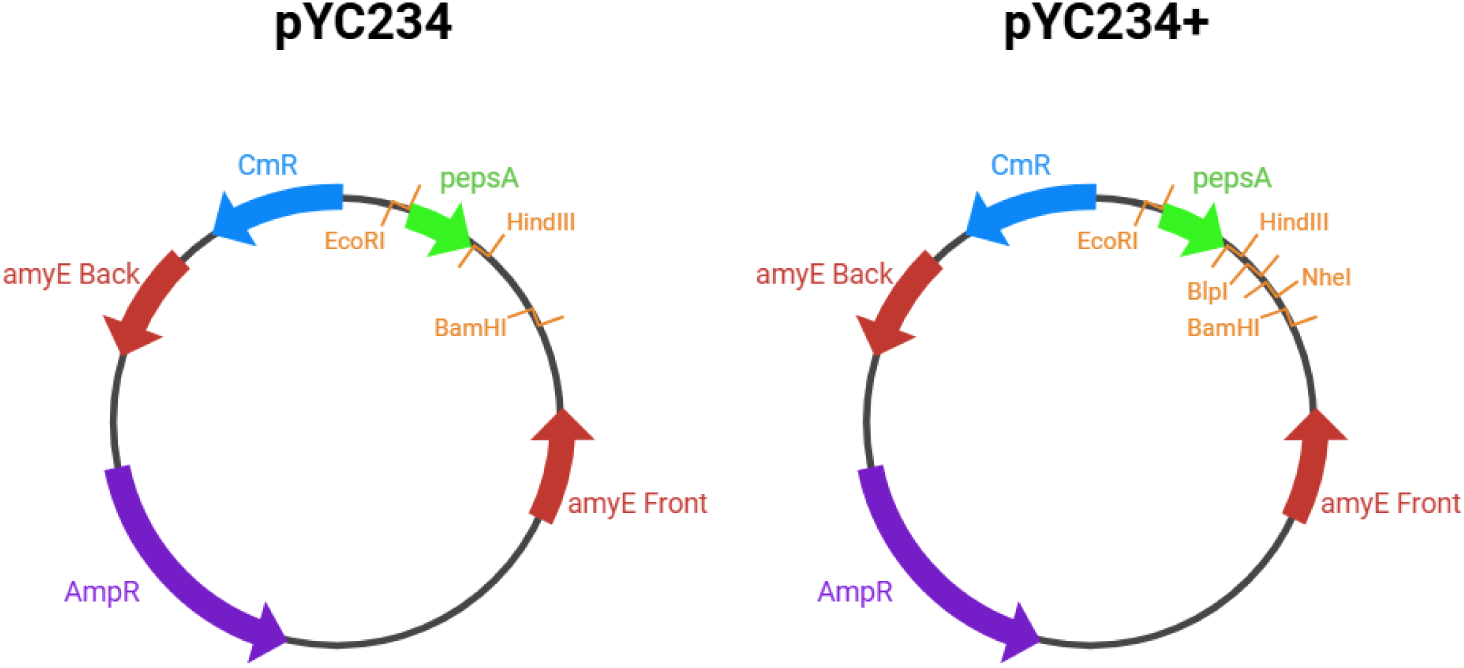
pYC234 and pYC234+ Vector Maps.

To facilitate the subsequent insertion of individual *eps* gene, additional restriction sites were introduced into pYC234 in the region between BamHI and HindIII sites. Primers pYC234+_Forward (5’-GTACGGATCCGCTAGCGCTAAGCAAGCTTATAGCCTTCAGCCTTC-3’) and pYC234+_Reverse (5’-CCGATGATAAGCTGTCAAAC-3’) were used to amplify the region of pYC234 between the BamHI and HindIII restriction sites while also adding NheI and BlpI sites between the existing sites using Gloria Nova HS 2X Master Mix (ABclonal, Woburn, MA, USA, 01801).

### Construction of *eps* gene complementation strains

A set of custom plasmids was generated using pYC234+; each plasmid harbors a different *eps* gene in the *eps* operon under the control of the *epsA* promoter. *eps* genes were amplified using the primers listed in Table 4 and Gloria Nova HS 2X Master Mix (ABclonal, Woburn, MA, USA, 01801). PCR products were purified using the GeneJET PCR Purification Kit (Thermo Fisher Scientific, Waltham, MA, USA). The *eps* genes and purified plasmid DNA of pYC234+ (purified using GeneJET Plasmid Miniprep Kit, Thermo Fisher Scientific, Waltham, MA, USA) were digested with the two restriction enzymes specified for each construct in Table 4 (New England Biolabs, Ipswich, MA, USA 01938). The digested genes and linearized pYC234+ were ligated together using T4 DNA ligase (ABclonal, Woburn, MA, USA, 01801) for 45 minutes at room temperature.

**Table 4:**
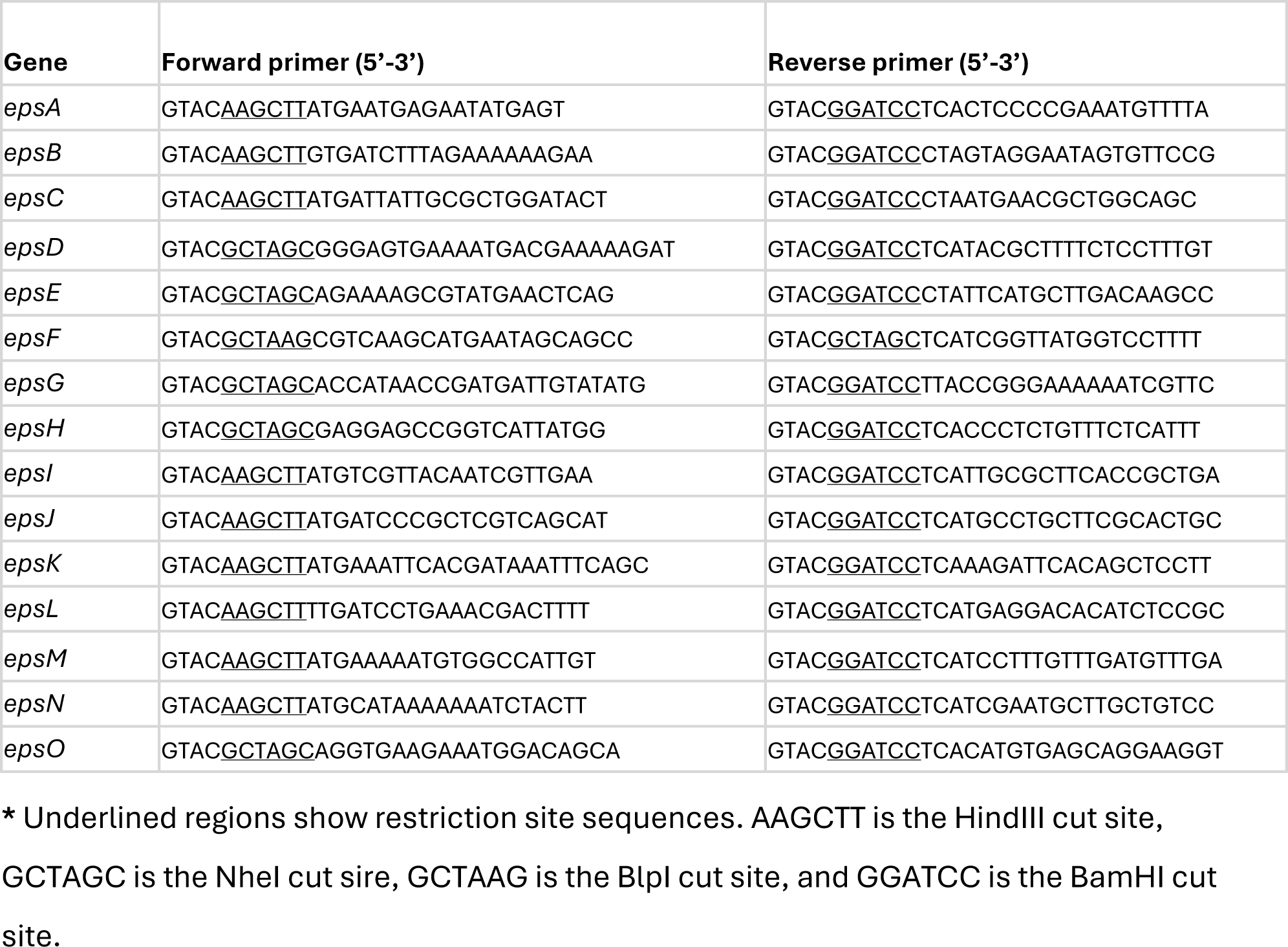
Complementation Strain Primers and Restriction Enzymes.

Each plasmid-gene ligation was transformed into chemically competent DH5α *E. coli*. Briefly, 50 µL of competent DH5α cells were mixed with 10 µL of purified plasmid on ice for 30 minutes. The cells were transferred to a 42°C water bath for 45 seconds (heat shock) and then returned to ice for 2 minutes. Following the addition of 1mL LB, the cells were incubated while shaking at 37°C for 1 hour. A 100 µL aliquot was then plated on LB agar containing 100 µg/mL ampicillin and incubated at 37°C overnight. Putative transformants were picked and screened by colony PCR using the indicated primers (Table 4). For positive clones, DNA sequencing was applied to verify the correct insert, and absence of PCR-introduced mutations.

Each pYC234+ plasmid containing an *eps* gene was transformed into *B. subtilis* strain PY79, where it recombined with the *B. subtilis* genome at the *amyE* site. PY79 was grown in MC medium for 4 hours with shaking at 37°C. A 300µL aliquot of this culture was combined with 10 µL of the purified plasmid. Cells and plasmids continued to incubate shaking at 37° C for an additional 2 hours. The entire volume was then plated on an LB plate containing 7 µg/mL chloramphenicol and grown overnight at 37°C. Putative transformant colonies were picked and re-streaked onto LB agar containing 7 µg/mL chloramphenicol and onto starch-containing agar plates. The plates were incubated overnight at 37°C.

Integration at the *amyE* locus was initially screened using the starch-hydrolysis assay. Starch plates were flooded with iodine solution, which stains unhydrolyzed starch dark blue or black. Because integration at *amyE* disrupts the endogenous amylase gene, correctly integrated colonies are expected to remain amylase-negative and therefore lack a clear halo. Colonies that have clear halos around them are amylase positive, and were considered to have retained an intact *amyE* locus. gDNA from colonies with successful gene integration was isolated using the Wizard Genomic DNA Purification Kit (Promega, USA) according to kit instructions. Extracted gDNA was then transformed into the *B. subtilis eps* gene deletion strains in NCIB 3610. The transformation was carried out as described above for transformation of plasmids into strain PY79. In this case, 10 µL of PY79 gDNA was used in place of purified plasmids.

### Biofilm development and imaging

One day prior to inoculation, fresh LB agar plates containing 1% (w/v) glycerol and 0.1 mM MgSO_4_ (LBGM, a biofilm inducing medium) were made [36]. The plates were allowed to dry for about 30 minutes before inoculation. Liquid cultures of each selected colony were grown in LB broth supplemented with 7 µg/mL chloramphenicol for 4 hours shaking at 37°C. For colony biofilm assay, 2 µL of each culture was spotted gently onto the agar.

For pellicle biofilm assay, 3 mL of LBGM media was aliquoted into 12 well culture plates. A 3 µL aliquot of each liquid culture was added to a well to obtain a 1:1000 dilution. The plates were gently shaken to mix. Following inoculation, all plates were incubated at 30°C under static conditions. Biofilms were imaged at 24, 48, and 72 hours after inoculation using Leica DMC2900 (Wetzlar, DEU) dissecting microscope at 2x to 8x magnification. Images were collected from biological replicates and from comparable location within each colony or pellicle.

## Results and Discussion

To assess the impact of *eps* gene deletion as well as complementation on biofilm phenotype, the rugosity and overall morphology of colony and pellicle biofilms were assessed. Strains that formed pellicle and colony biofilms with highly wrinkled phenotypes were considered robust biofilm formers. Biofilms that displayed mildly wrinkled or textured morphologies were considered moderate or altered biofilm formers, whereas biofilms that lacked wrinkled morphology were considered biofilm-deficient. In biofilm deficient strains, mucoid colonies typically formed on agar surfaces, and cells typically settle at the bottom of standing liquid cultures instead of forming floating pellicles. These phenotypic categories were assigned based on visual comparison with the wild-type NCIB 3610.

Based on the observed biofilm phenotypes, each gene in the *eps* operon contributed to robust biofilm (Figure 4). Biofilms formed by the deletion strains differ from those by the wild-type NCIB 3610, leading to biofilm-deficient or occasionally altered phenotypes. The absence of some genes, including *epsA*, *epsB*, *epsJ*, and *epsO*, still allowed formation of a moderate biofilm (Figure 4)*. epsA* and *epsB* encode regulatory proteins for EPS chain length control (Table 5). *epsJ* encodes a glycosyltransferase predicted to add a branching galactose residue onto the EPS glycan (Table 5). *epsO* encodes a predicted pyruvyl-transferase that adds a pyruvyl group to the branching galactose (Table 5).

**Figure 4:**
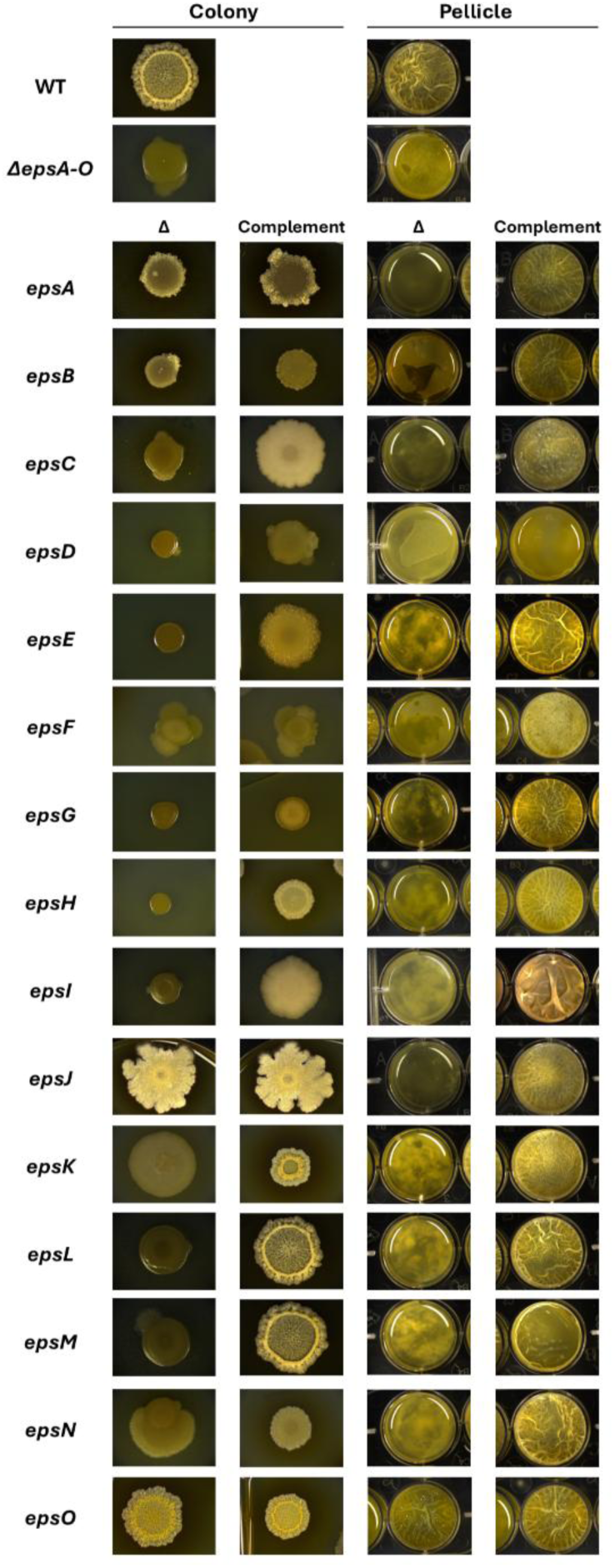
Colony and pellicle biofilms grown from *eps* non-polar deletion and complementation strains in an NCIB 3610 background. All single gene deletions alter biofilm phenotype compared to the wild type.

**Table 5:**
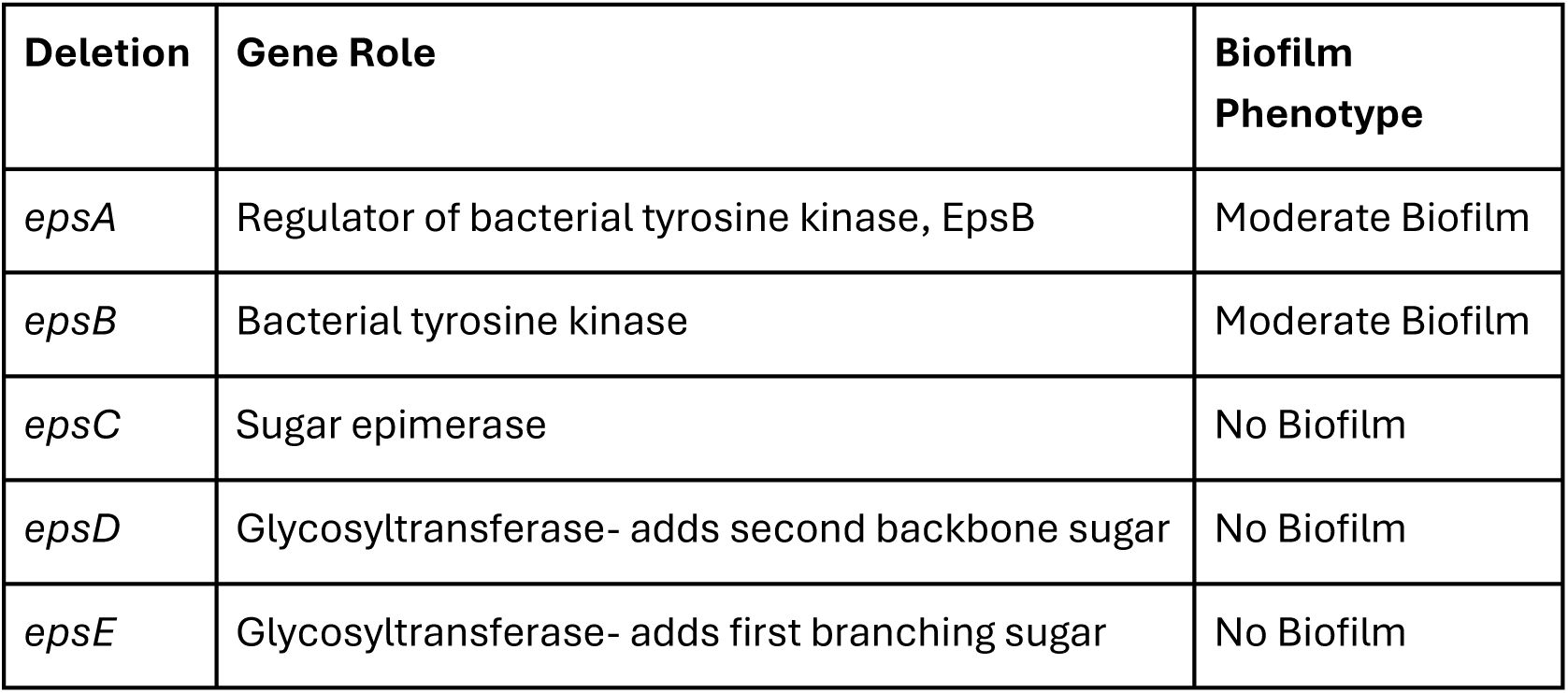
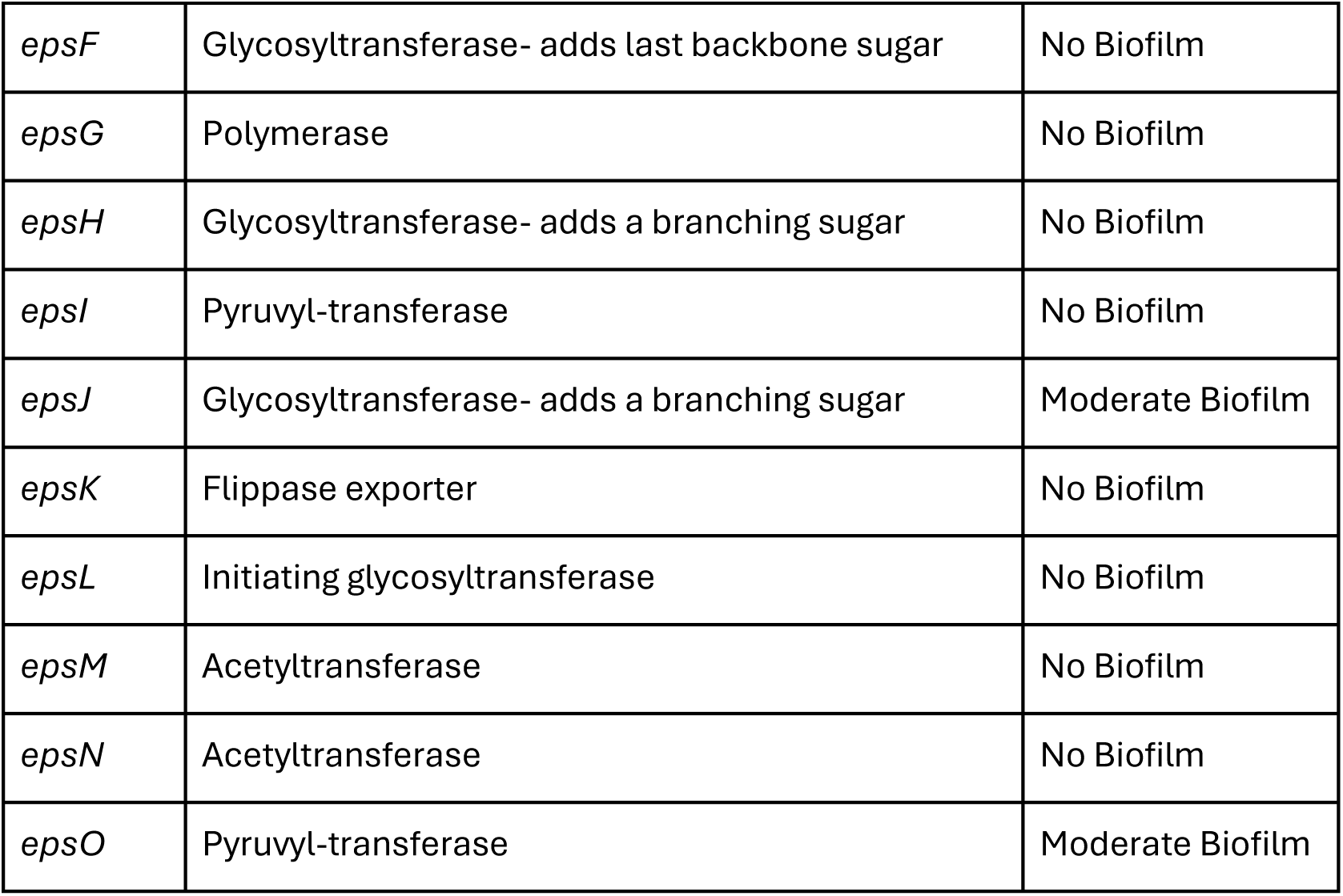
Summary of biofilm phenotypes in marker-less, in frame deletion strains.

For each gene deletion, complementation of the respective gene driven by the *epsA* promoter at the *amyE* site restores wild type-like biofilm growth phenotype, or at least partially (Figure 4). This confirms that the marker-less in-frame gene deletions made are non-polar, and gene absence does not impact the expression of downstream genes.

Based on the results of this study, all genes in the *eps* operon seem to be required for robust biofilm development. However, four of the fifteen *eps* genes, *epsA*, *epsB*, *epsJ*, and *epsO*, are not strictly essential since their loss still enables a moderate biofilm phenotype. The absence of *epsA* or *epsB* still allows for the formation of an altered biofilm phenotype in both colony and pellicle biofilm settings (Figure 4). This was not surprising since their functions are regulatory and neither gene contributes directly to EPS biosynthesis. Intriguingly, the absence of two other genes that play direct roles in EPS biosynthesis, *epsJ* or *epsO,* still allows biofilm formation, albeit with moderately impaired phenotypes (Figure 4). The very moderate biofilm phenotype of Δ*epsJ* was especially surprising. It was originally hypothesized that the deletion of any GT gene would be fully detrimental to biofilm formation, since they play central roles in defining overall sugar structure. Δ*epsO* that has a partial phenotype was also surprising. While it hasn’t been deeply explored in *B. subtilis*, pyruvyl groups have been shown to serve key functions for polysaccharides, including cross-linking with proteins [37]. Looking at the structure of EPS proposed by Dogsa et al. in 2024 helps shed light onto why these seemingly important genes are not essential for biofilm formation as we further discuss below.

Mature EPS molecules generally contain six total sugars-three backbone sugars and three branching galactose molecules that branch off of the center backbone molecule [13]. However, it was found that a subset of purified EPS molecules only has two sidechain galactose molecules rather than three. This suggests that the last galactose on the side chain may be conditional or optional, not critically important for assembly of a robust biofilm. Rather, it may be beneficial for biofilm stability. Both EpsH and EpsJ are predicted galactose-transferases. It was unclear which enzyme is responsible for adding the final galactose molecule onto the sidechain. Based on sequence homology alone, it is not possible to determine if EpsH or EpsJ is responsible. In this study, Δ*epsH* is incapable of forming biofilms. In fact, Δ*epsH* has long been used as a biofilm deficient strain and a negative control in *B. subtilis* [38]. Δ*epsJ* on the other hand forms a moderate biofilm.

Looking at the structural data and this biofilm growth phenotype together, it appears that EpsJ may be responsible for adding the final side-chain galactose to the molecule (Figure 5), while EpsH may add the middle galactose to the side-chain.

A similar deduction can be made regarding the functions of EpsI and EpsO. Both enzymes are predicted to be responsible for pyruvylating the two side-chain galactose molecules. Given that Δ*epsI* forms severely defective biofilms while *ΔepsO* develops a moderate biofilm phenotype, this could be the evidence that EpsI is responsible for modifying the terminal galactose sugar in the side chain, while EpsO is more likely to modify the middle branching galactose molecule, and both that middle galactose and the pyruvyl-modification are critically important for robust biofilm development (Figure 5).

When performing assessments that rely heavily on biofilm phenotype analysis, it was important to grow biofilms both in colony- and pellicle-forming conditions. In this study, moderate biofilm growth phenotypes are more apparent in colonies grown on solid medium than in pellicles grown in liquid culture (Figure 4). Having the support of a solid surface helps showcase mild biofilm phenotypes that may be lost because they struggle to form on a less supportive liquid surface. Furthermore, successful complements of certain gene deletions, especially those which had partial biofilm phenotypes to begin with, were more apparent in pellicle biofilms.

This method of using single-gene deletion strains to help elucidate the specific roles of genes in an operon is widely adaptable and can be applied to a variety of operons and strains of bacteria. There are limitations to this method. The order in which the GTs in the *eps* operon act could be further confirmed through Nuclear Magnetic Resonance (NMR) of EPS purified from different GT deletion strains. When combined with biochemical data, it can be a powerful and relatively low-cost tool for helping infer gene functions.

## Conclusion

Single-gene deletion mutants are a useful tool for elucidating the roles of biosynthetic genes in large operons. For example, even though the structure of EPS was identified in 2024, it was not possible to differentiate and determine the roles of EpsH and EpsJ through sequence-based homology alone [13]. Through the development of a single-*eps* gene deletion library in wild type *B. subtilis*, we propose that EpsJ is the last-acting GT on the EPS 3-ring side chain as its absence was not detrimental to biofilm formation.

Similarly, we propose that EpsO pyruvylates the last side chain sugar of EPS while EpsI pyruvylates the second side chain sugar. This work brings us one step closer to establishing the *eps* operon as a model for complex exopolysaccharide formation in Gram positive organisms.

## Acknowledgements

We thank the National Science Foundation (NSF) for supporting this work (MCB1651732). The NSF also supported N.T.C. through the Graduate Research Fellowship Program (GRFP) (1938052). N.T.C was supported through the PhD Network’s Dissertation Completion Fellowship at NU. B.P.U was supported through the AJC Merit Research Scholars program at NU. E.V.F. and E.W. were funded by Project-Based Exploration for the Advancement of Knowledge (PEAK) fellowships at NU.

## Notes

### Competing Interest Statement

The authors have declared no competing interest.

